# Neonatal Muscle Tone Predicts Cerebellar Morphology Later in Development Without Mediating Autistic Traits

**DOI:** 10.64898/2026.08.24.746825

**Authors:** Dylan van der Waal, Amina Z. Z. Burgess, Wietske van der Zwaag, Aleksandra Badura, Bing Xu, Serena Defina, Alexander Neumann, Pauline W. Jansen, Ryan Muetzel, Carolin Gaiser

## Abstract

**Background:** Infant muscle tone reflects early central nervous system integrity and has been associated with later motor and cognitive development, including autism traits. The cerebellum regulates both motor control and higher-order socio-cognitive functions and has been repeatedly implicated in autism, but its role in linking infant muscle tone to adolescent autistic traits has not previously been studied in a large, prospective population cohort.

**Methods:** We used data from the prospective Generation R Study. Infant muscle tone (hypotonia and hypertonia) was assessed via Prechtl examination, and third-trimester fetal transcerebellar diameter was measured using ultrasound *n*=6,842). Cerebellar morphology at ages 6, 10, and 14 years (*n*=4,861) was measured using structural MRI. Linear mixed-effects models tested associations between infant muscle tone and 35 anatomical and 10 functional cerebellar regions. Causal mediation models tested whether cerebellar volume mediated associations between infant muscle tone and adolescent autistic traits at age 14 (Social Responsiveness Scale).

**Results:** Hypotonia predicted larger vermis IX volumes across childhood (*beta*=0.037, *pFDR* =0.043). Hypertonia showed an age-dependent association with left lateral lobule IX (*beta*=−0.0027, *pFDR* =0.041), with differences diminishing with age. Third-trimester transcerebellar diameter did not predict postnatal muscle tone. Given its significant main effect, vermis IX volume was tested as a mediator, but did not mediate the pathway to adolescent autistic traits. However, infant hypotonia showed a small direct association with elevated autistic traits at age 14, specific to girls (*beta*=0.0255, *p*=0.020).

**Conclusions:** Infant muscle tone is associated with localized differences in cerebellar volumes. These associations are specific to vermal and left hemispheric lobule IX, a region commonly implicated in spinocerebellar postural control, axial stability, and higher-order sensorimotor integration. Furthermore, infant muscle tone was not predicted by prenatal cerebellar diameter, and cerebellar volumes did not mediate the association between infant hypotonia and adolescent autistic traits in our study. Future research should further investigate these findings in clinical populations, integrating longitudinal whole-brain, multi-modal imaging to clarify the association between infant muscle tone, the cerebellar functioning, and autistic traits.

## Background

Infant muscle tone is a primary clinical indicator of central nervous system maturation, offering an early observational window into neurodevelopmental trajectories (Bhat, 2020; Ganguly et al., 2021; Lance, 1980; Serdarevic et al., 2017; Spittle et al., 2008). Muscle tone sensitively reflects network integrity, acting as a functional output that depends on seamless coordination across the cortex, cerebellum, brainstem, spinal cord, and muscle spindles (Dow, 1961; Gilman, 1969; Martin, 2021; Pierrot-Deseilligny & Burke, 2005). Disruptions of these circuits typically present as either hypotonia, characterized by decreased resistance to passive stretch, or hypertonia, defined by excessive stiffness (Campbell & Barohn, 2019; Lance, 1980). Rather than representing isolated motor deficits, early muscle tone deviations, particularly hypotonia, serve as sensitive markers of atypical neurodevelopment. Observational motor evaluations like Prechtl’s assessment even outperform early MRI in predicting cerebral palsy (Novak et al., 2017), while also marking elevated risk for autism spectrum disorder (ASD) (Bhat, 2020; Serdarevic et al., 2017).

Beyond these early motor indicators, ASD is characterized by persistent impairments in social communication and restrictive, repetitive behaviors, with subclinical autistic traits extending continuously throughout the general population (American Psychiatric Association, 2013; Constantino & Todd, 2003). These socio-communicative and behavioral features are increasingly understood to stem from early, systemic alterations in brain network connectivity (Stoodley et al., 2017; Wang et al., 2014), directly linking early neuromotor vulnerabilities with developing social-cognitive outcomes (Bhat, 2020; Licari et al., 2020). While motor deviations manifest earliest in infancy, higher-order socio-communicative features emerge later during development (Bhat, 2020; Serdarevic et al., 2017). Understanding which neural structures could plausibly link early muscle tone deviations to ASD is therefore essential for understanding early developmental pathways (Ganguly et al., 2021; Stoodley & Schmahmann, 2009).

The cerebellum might be a promising candidate for this link. Beyond its established role in motor regulation, structural neuroimaging consistently identifies the cerebellum as one of the most altered brain regions in autistic individuals (Fatemi et al., 2012; Stoodley et al., 2017).

Functionally, cerebellar outputs directly tune the descending loops responsible for muscle tone, while its subregional functional topography covers features across motor, cognitive, and affective domains (Buckner et al., 2011; Strick et al., 2009). Undergoing rapid cellular growth during the third trimester of pregnancy and early infancy, the cerebellum is exceptionally vulnerable to early developmental disruptions (Choe et al., 2013; Wang et al., 2014). While early work established that the cerebellum continues to mature into adolescence (Tiemeier et al., 2010), recent population-wide normative modeling revealed an anterior-to-posterior growth gradient, with motor areas maturing earlier than posterior cognitive subregions (Gaiser et al., 2024). Topographically, anterior lobules (I–VI) along with posterior lobules VIII and IX support sensorimotor control and posture, whereas posterior lobules Crus I, Crus II, and VIIB support higher-order socio-cognitive processing. However, despite extensive evidence independently implicating the cerebellum in muscle tone regulation and ASD neurobiology, these interconnected pathways have not been evaluated within a combined longitudinal framework (Kelly et al., 2020; Stoodley et al., 2017).

In this study, we used prospective longitudinal data from the Generation R cohort, to map cerebellar development from fetal life to adolescence, linking early infant muscle tone deviations to later cerebellar morphology and autistic traits. First, we examined whether infant muscle tone deviations predict subregional cerebellar morphology across childhood and early adolescence. Second, to probe the directionality of this relationship, we tested whether prenatal cerebellar size predicts infant muscle tone. Third, we examined whether cerebellar morphology could potentially mediate the relationship between infant muscle tone and autistic traits in adolescence. Together, these analyses shed light on the neurodevelopmental links between early motor development, cerebellar structure, and autistic traits, with implications for understanding how early neuromotor features contribute to the etiology of autism.

## Methods and Materials

### Participants

This study was embedded within the Generation R Study, a population-based prospective cohort in Rotterdam, the Netherlands, which follows children from fetal life into young adulthood. Pregnant women with an expected delivery date between April 2002 and January 2006 were recruited (N = 9,778) (Kooijman et al, 2016). The study has been approved by the Medical Ethical Committee of the Erasmus MC, University Medical Centre in Rotterdam. Participants did not receive monetary compensation, but their travel costs were reimbursed. Additionally, as a token of appreciation for their participation, they received small gifts valued at 10€ or less. Written informed consent was obtained from all participants. The current study employs an observational, mixed longitudinal design using data collected during the prenatal period, infancy, and at ages 6, 10, and 14 years.

### Measurements

#### Infant Neuromotor Assessment

Infant muscle tone was evaluated as part of the early neuromotor development assessment between 8- and 27-weeks post-term age using the Prechtl Neurological Examination (Prechtl, 1977). Tone was examined across multiple postures (supine, horizontal suspension, vertical suspension, prone, and sitting) and specific clinical items (e.g., adductor angle), with each item scored as normal, low, or high tone (Supplementary Table 1). Corrected sum scores were computed separately for low tone (hypotonia) and high tone (hypertonia) resulting in two continuous variables.

#### Autistic Traits

Autistic traits were assessed at age 14 using an 18-item short form of the Social Responsiveness Scale (SRS; Constantino & Gruber, 2005). The SRS is a parent-report questionnaire measuring quantitative subclinical and clinical autistic traits over the preceding six months across both DSM-5 ASD domains: social communication/interaction and restricted, repetitive patterns of behavior (Constantino & Todd, 2003). Each item is scored on a 4-point scale ranging from 0 (“never true”) to 3 (“almost always true”), with higher scores indicating greater endorsement of autistic traits. To minimize participant burden, an 18-item abbreviated version of the SRS was administered (Roman et al., 2013). This short form demonstrates high construct validity with the full 65-item SRS, showing high correlations in both Dutch clinical/population-based validation samples (r = 0.95) and independent twin cohorts (r = 0.93 - 0.94) (Blanken et al., 2015; Constantino & Todd, 2003). Weighted total scores were computed based on non-missing items, excluding individuals with more than 5 missing items.

#### Prenatal Transcerebellar Diameter

Fetal ultrasound examinations were performed by qualified sonographers across all three trimesters. Most assessments (88%) were conducted at the central Generation R Research Centre in Rotterdam, with the remainder carried out at five collaborating regional hospitals. These ultrasounds were used to determine gestational age and track fetal growth. The transcerebellar diameter (TCD) was measured in millimeters across the widest axial plane of both cerebellar hemispheres using a slightly caudal-rotated view. To address our secondary research question, whether prenatal cerebellar size predicts infant muscle tone, we analyzed transcerebellar diameter measured during the third trimester (Between 25 and 40 weeks of gestation). We specifically selected third-trimester measurements to capture the period of rapid cerebellar growth (Choe et al., 2013; Wang et al., 2014).

#### Cerebellar Neuroimaging

##### Image Acquisition and Preprocessing

Structural T1-weighted magnetic resonance imaging (MRI) was performed in three prospective data collection waves at mean ages 6, 10, and 14 years. Scans were acquired on two 3T GE MRI systems: the initial assessment used a GE MR750 Discovery scanner, while all subsequent assessments were conducted on a study-dedicated GE MR750w system (General Electric Healthcare, Milwaukee, WI, USA). High-resolution structural images were acquired using inversion recovery fast spoiled gradient-recalled sequences (IR-FSPGR) (White et al., 2013).

Structural preprocessing followed standard pipelines via SMRIPrep (Esteban et al., 2021). Briefly, scans at age 6 were resampled to 1 mm³ isotropic resolution to align spatially with subsequent waves. Preprocessing included skull-stripping, B1 field inhomogeneity correction, and spatial registration to MNI152 template space (NLin2009cAsym, 1 mm³ resolution) using the Advanced Normalization Tools (ANTs). We calculated the Jacobian determinant (measure of volume of a given voxel relative to its volume in MNI152 space) from the nonlinear warp field. Tissue segmentation was performed with FSL FAST to yield voxel-wise probability maps of grey matter densities (GMD), white matter densities (WMD), and cerebrospinal fluid (CSF). All scans were visually inspected to ensure quality.

##### Cerebellar Atlases and Regional Extraction

To capture both structural and functional cerebellar properties, two complementary parcellation atlases were used. For anatomical parcellation, the Multiple Automatically Generated Templates (MAGeT Brain) algorithm (Park et al., 2014) was used to extract 35 anatomical regions of interest (ROIs), encompassing individual cerebellar lobules across both hemispheres, vermal subregions, and the bilateral corpus medullare. For functional parcellation, the Multi-Domain Task Battery (MDTB) atlas (King et al., 2019) was applied to segment the cerebellum into 10 functional regions based on cognitive, motor, and socio-emotional task activation patterns. Following the functional MDTB atlas of King et al. (2019), these 10 functional regions contain Left-hand motor (Region 1), Right-hand motor (Region 2), Saccades/Vermis (Region 3), Action observation (Region 4), Divided attention (Region 5), Active maintenance (Region 6), Narrative/Language (Region 7), Word comprehension (Region 8), Verbal fluency (Region 9), and Autobiographical recall (Region 10).

##### Normative Modeling

To account for the scanner differences between the visit at age 6 and subsequent visits normative modeling was applied using the PCNtoolkit Python package (v1.2.0; Python v3.10.6) (Marquand et al., 2016; Rutherford et al., 2022; de Boer et al., 2024; Kia et al., 2022). Normative models for each anatomical (35 ROIs) and functional (10 ROIs) cerebellar subregion were estimated as a function of age, controlling for batch effects of sex and scanner. To evaluate model stability and prevent overfitting, the dataset was split into 50% training and 50% testing sets, stratified by sex and scanner. This cross-validation split was repeated across five random iterations, reversing training and testing sets in each iteration to generate batch-effect-free normative estimates across the entire cohort. Stability across the five resampled trainings fold iterations was excellent, with Intraclass Correlation Coefficients (ICCs) ranging from 0.988 to 0.999 across 35 anatomical ROIs and from 0.993 to 0.999 across 10 functional ROIs. Normative estimates across the five iterations were therefore averaged, resulting in a single normative estimate per ROI and participant.

##### Covariates

All statistical models were adjusted for the following potential confounders: biological sex, gestational age at birth, parental national origin, household income, maternal education level, third-trimester maternal blood pressure (systolic and diastolic), and prenatal substance use (smoking, alcohol, and benzodiazepine use). Models evaluating MRI-based outcomes were additionally adjusted for age at scan and intracranial volume (ICV). Biological sex and gestational age at birth were obtained from midwife and hospital registries at birth. National origin of the parents was categorized into three distinct categories based on parental national origin (1 = Dutch, 2 = Non-Dutch, European, 3 = non-European). Household income, defined as the total net monthly income of the household, was categorized into <1200 Euro (“low”), 1,200– 3,200 Euros (“middle”), and >3,200 Euros (“high”). The education level of the mother and father was assessed by the highest completed education and reclassified into two categories: “low” (at most intermediate vocational training), and “high” (at least college education). Maternal smoking, alcohol use, and benzodiazepines use during pregnancy were simplified into two categories (Yes/No). Maternal blood pressure was measured during the third trimester ultrasound visit at the Generation R Research Centre.

Missing covariate data were imputed across 30 datasets (m=30, 40 iterations) using parallelized multiple imputation by chained equations (futuremice in R). Variable-specific algorithms were applied based on measurement scale: random forest (ranger) for continuous metrics, logistic regression for binary indicators, and polytomous logistic regression for unordered categorical variables.

### Statistical Analysis

#### Main analysis

To examine the longitudinal association between infant muscle tone and longitudinal cerebellar development, separate Linear Mixed-Effects Models (LMM) were fitted for hypotonia and hypertonia. The main predictors were infant muscle tone sum scores (hypotonia or hypertonia), and the outcomes were cerebellar volumes across 45 ROIs (35 anatomical + 10 functional ROIs). To model the repeated MRI measurements across the three time points (ages 6, 10, and 14 years), a random intercept per participant was specified. The full statistical models were defined as follows:

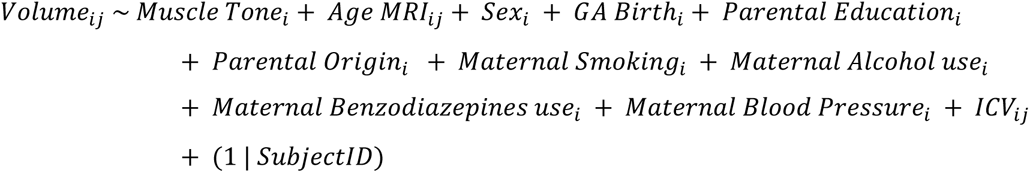

*i indexes the individual subject (i=1,…,N)*.

*j indexes the specific scan or time point for subject i*.

To address selection bias and non-random participation across follow-up waves, Inverse Probability Weighting (IPW) was applied. Using a stepwise logistic regression approach, a composite weight was calculated by multiplying the inverse probability of inclusion at age 6 (conditional on baseline parental and demographic covariates) by the conditional probabilities of inclusion at ages 10 and 14 (conditional on participation at age 6). These composite weights were incorporated into all analyses involving MRI data (Dijkzeul et al., 2024). Multiple testing was controlled using the False Discovery Rate (FDR) correction.

#### Secondary Analyses

Two secondary analyses were conducted to clarify directionality and potential behavioral mechanisms. First, to test directionality, we tested whether prenatal transcerebellar diameter assessed via third-trimester fetal ultrasound predicted infant muscle tone (hypotonia or hypertonia). Because the infant muscle tone scores are zero-inflated due to a high proportion of neurodevelopmentally typical scores, hurdle regression models were implemented (Mabire-Yon, 2025). These models were adjusted for gestational age at ultrasound scan alongside the prenatal and sociodemographic covariates, excluding post-natal MRI-specific metrics.

Second, causal mediation analysis was performed to evaluate whether longitudinal cerebellar morphology mediated the association between infant muscle tone and adolescent autistic traits at age 14. The SRS scores are square-root transformed total scores to handle the non-normal distribution. Longitudinal subregional metrics were summarized into individual developmental trajectory parameters using linear mixed-effects modeling to extract adjusted participant-level values across the three waves (See Figure 1).

**Figure 1.**
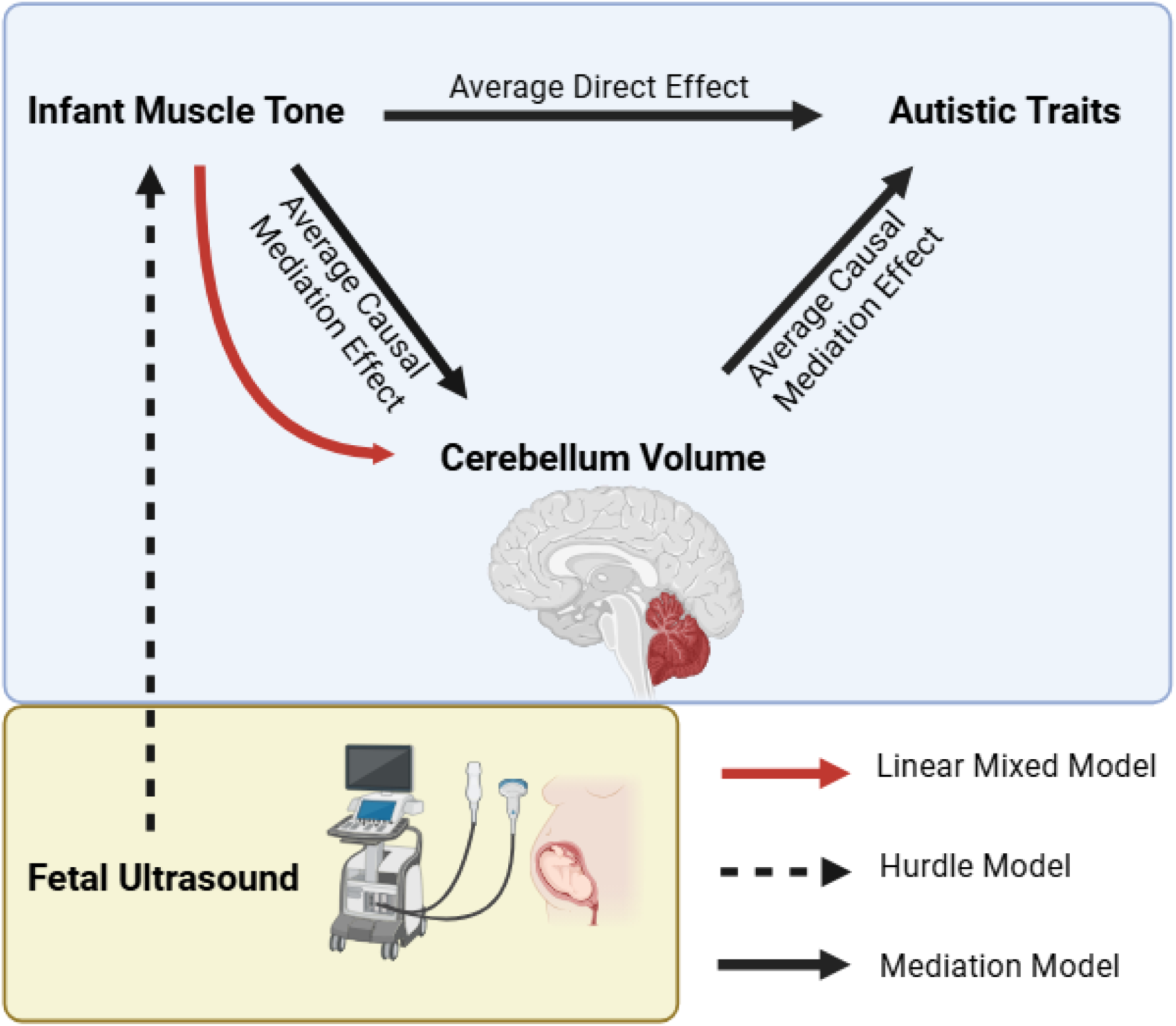
Overview of Statistical Models. *Note.* Primary analysis evaluates longitudinal trajectories between infant muscle tone and cerebellar volume (red). Secondary analyses test prenatal ultrasound predicting infant muscle tone via hurdle models (dashed black) and causal mediation to adolescent autistic traits (solid black).

Causal mediation was modeled within a linear regression framework using the mediation R package to estimate the average causal mediation effect (ACME), average direct effect (ADE), total effect, and proportion mediated. Models were adjusted for the full set of sociodemographic, maternal, and imaging covariates. Unbiased non-parametric bootstrap confidence intervals were derived using 3,000 simulations. To investigate sex-specific neurodevelopmental pathways, mediation analyses were conducted in the total cohort and subsequently stratified by sex.

## Results

### Participant Characteristics and Sample Selection

From the initial 9,778 pregnant women enrolled into the Generation R Study, 7,893 consented to postnatal follow-up. A total of 5,185 participants completed structural MRI neuroimaging across ages 6, 10, or 14 years, of whom 4,861 had usable MRI data following quality control checks (Figure 2).

**Figure 2.**
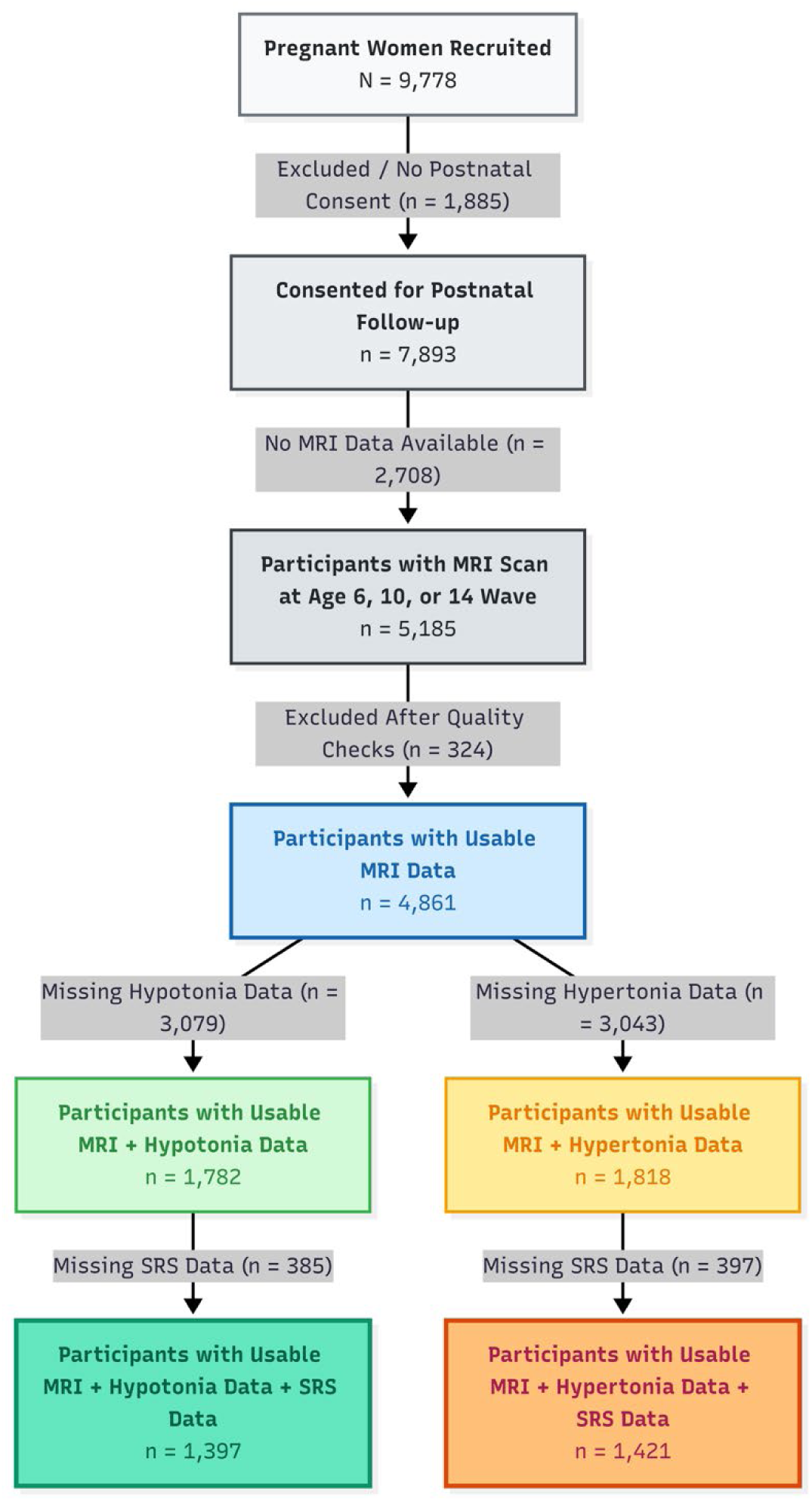
Flowchart of Study Population and Analytical Subsample Derivation. *Note.* Overview of participant inclusion from the baseline Generation R pregnant cohort through quality control checks, longitudinal neuroimaging waves (ages 6, 10, and 14 years), infant muscle tone analytical tracks (hypotonia and hypertonia), and final behavioral mediation samples. Abbreviations: MRI = Magnetic Resonance Imaging; SRS = Social Responsiveness Scale.

For the primary analyses, investigating whether muscle tone in infancy predicts cerebellar development, two non-mutually exclusive samples were derived based on the availability of infant muscle tone assessments: a hypotonia sample (n=1,782; 2,679 total usable MRI scans) and a hypertonia sample (n=1,818; 2,732 total usable MRI scans) (Figure 2).

Demographic profiles were highly similar across both muscle tone samples (Table 2a, 2b). The hypotonia sample comprised 923 females (51.8%) and 859 males (48.2%), with an ethnic distribution of 55.4% Dutch ancestry, 9.9% non-Dutch European ancestry, and 34.7% (n=610) non-European ancestry. Similarly, the hypertonia sample comprised 942 females (51.8%) and 876 males (48.2%), with 55.1% of Dutch ancestry, 9.9% non-Dutch European ancestry, and 34.9% (n=627) non-European ancestry. Maternal educational level, household net income, and maternal substance use during pregnancy (smoking, alcohol, and benzodiazepines) showed matching distributions between groups (Table 2a, Table 2b, Figure 3).

**Figure 3.**
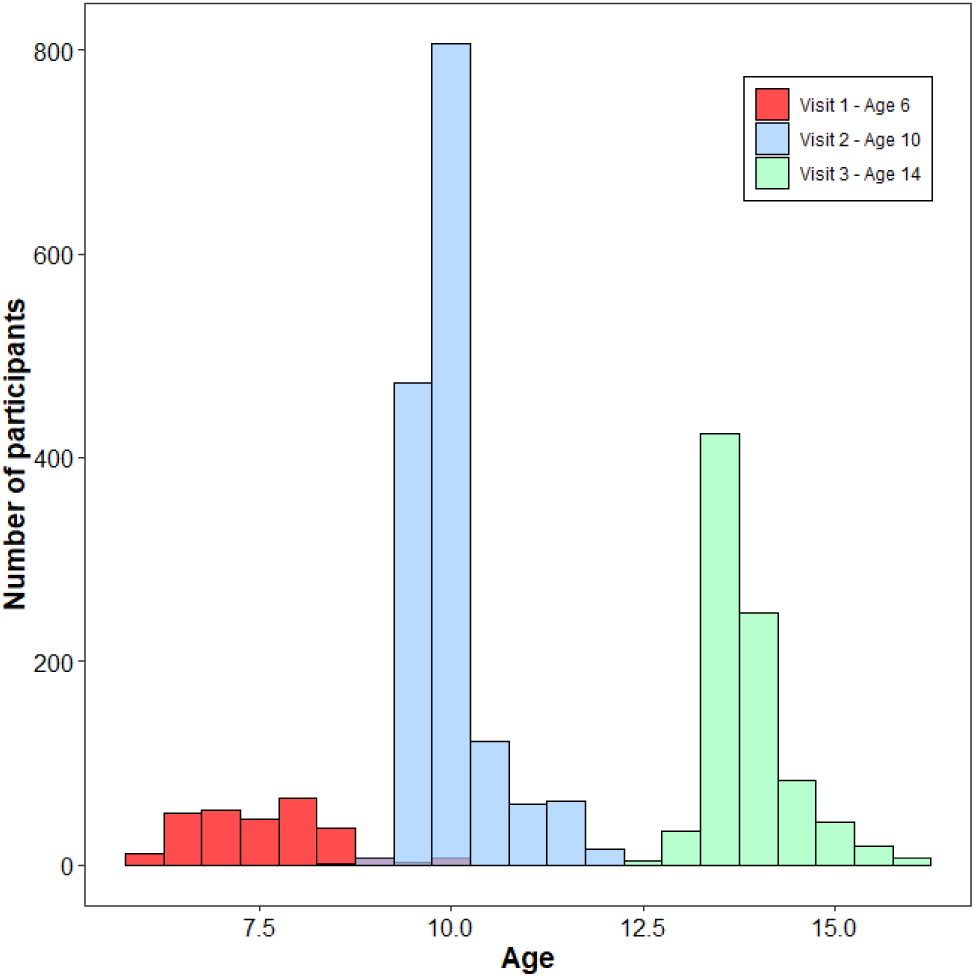
Age and Scanner Distribution Across Assessment Waves for the Muscle Tone Analytical Subsamples. *Note.* Density distributions of participant age (in years) and scanner model allocation across the three longitudinal data collection waves (ages 6, 10, and 14 years) for the hypotonia (n=1,782) and hypertonia (n=1,818) cohorts. -axis represents the total number of usable structural MRI-scans per assessment bin.

**Table 2a.** a Participant Characteristics of Hypotonia Subgroup.

|  | *Level* | *Overall* | *Missingness (%)* |
| --- | --- | --- | --- |
| Child Characteristics (*N*) |  | 1782 |  |
| Gestational age at birth (Mean (SD)) |  | 39.91 (1.71) | 0.1 |
| Sex (%) | Male | 859 (48.2) |  |
|  | Female | 923 (51.8) |  |
| Ethnicity (%) | Dutch | 974 (55.4) | 1.3 |
|  | Non-Dutch European | 174 (9.9) |  |
|  | Non-European | 610 (34.7) |  |
| Household Income (%) | < 1200 € | 94 (6.4) | 17.2 |
|  | 1200 – 3200 € | 580 (39.3) |  |
|  | > 3200 € | 801 (54.3) |  |
| Maternal Characteristics |  |  |  |
| Education (%) | Low | 775 (46.0) | 5.6 |
|  | High | 908 (54.0) |  |
| Alcohol use during pregnancy (%) | No | 614 (39.6) | 12.9 |
|  | Yes | 938 (60.4) |  |
| Smoking during pregnancy (%) | No | 1219 (77.9) | 12.2 |
|  | Yes | 345 (22.1) |  |
| Benzodiazepine use during pregnancy (%) | No | 1537 (99.0) | 12.9 |
|  | Yes | 16 (1.0) |  |
| Diastolic blood pressure during pregnancy (third trimester) (Mean (SD)) |  | 68.74 (8.94) | 8.7% |
| Systolic blood pressure during pregnancy (third trimester) (Mean (SD)) |  | 117.69 (11.31) | 8.7% |
| Paternal Characteristics |  |  |  |
| Education (%) | Low | 506 (43.3) | 34.5 |
|  | High | 662 (56.7) |  |

*Note.* *N* = Sample size, *SD* = Standard Deviation. Missingness % before imputation.

**Table 2b.** Participant Characteristics of Hypertonia Subgroup.

|  | *Level* | *Overall* | *Missingness (%)* |
| --- | --- | --- | --- |
| Child Characteristics (*N*) |  | 1818 |  |
| Gestational age at birth (Mean (SD)) |  | 39.88 (1.73) | 0.1 |
| Sex (%) | Male | 876 (48.2) |  |
|  | Female | 942 (51.8) |  |
| Ethnicity (%) | Dutch | 989 (55.1) | 1.3 |
|  | Non-Dutch European | 178 (10.0) |  |
|  | Non-European | 627 (34.9) |  |
| Household Income (%) | < 1200 € | 98 (6.5) | 17.2 |
|  | 1200 – 3200 € | 593 (39.4) |  |
|  | > 3200 € | 815 (54.1) |  |
| Maternal Characteristics |  |  |  |
| Education (%) | Low | 792 (46.2) | 5.6 |
|  | High | 924 (53.8) |  |
| Alcohol use during pregnancy (%) | No | 628 (39.7) | 12.9 |
|  | Yes | 955 (60.3) |  |
| Smoking during pregnancy (%) | No | 1241 (78.0) | 12.4 |
|  | Yes | 351 (22.0) |  |
| Benzodiazepine use during pregnancy (%) | No | 1570 (99.0) | 12.8 |
|  | Yes | 16 (1.0) |  |
| Diastolic blood pressure during pregnancy (third trimester) (Mean (SD)) |  | 68.73 (8.94) | 8.6% |
| Systolic blood pressure during pregnancy (third trimester) (Mean (SD)) |  | 117.66 (11.36) | 8.6% |
| Paternal Characteristics |  |  |  |
| Education (%) | Low | 516 (43.3) | 34.4 |
|  | High | 676 (56.7) |  |

*Note.* *N* = Sample size, *SD* = Standard Deviation. Missingness % before imputation.

For secondary analyses, third-trimester fetal ultrasound data were available in n=6,842 participants of which n=2,389 participants have hypotonia and n=2,427 participants have hypertonia assessments available. For adolescent causal mediation models at age 14, complete SRS behavioral data were available for n=1,397 participants in the hypotonia branch and n=1,421 in the hypertonia branch (Figure 2).

### Primary Analysis/Longitudinal Cerebellar Trajectories Associated with Infant Muscle Tone

#### Infant Hypotonia

After controlling for multiple testing across all models, a significant main effect of infant hypotonia was observed in Vermis IX (*β*=0.037, *SE*=0.015, *p*=0.013, *pFDR*=0.043, *95% CI* [0.008, 0.066]; Table 3, Figure 4A). Infants with higher hypotonia scores demonstrated larger volumes within this vermal region over time. Similar effects were observed in neighboring cerebellar regions, including Vermis VI (*p*=0.027), Vermis X (*p*=0.021), and left lobule VIIIB (*p*=0.044), which demonstrated larger volume estimates, however these regions did not survive FDR correction (*pFDR*>0.05; Table 3). Additionally, no significant associations were observed across the functionally defined ROI’s after FDR correction (*pFDR*>0.05; supplement figure 1 and 2)

**Figure 4.**
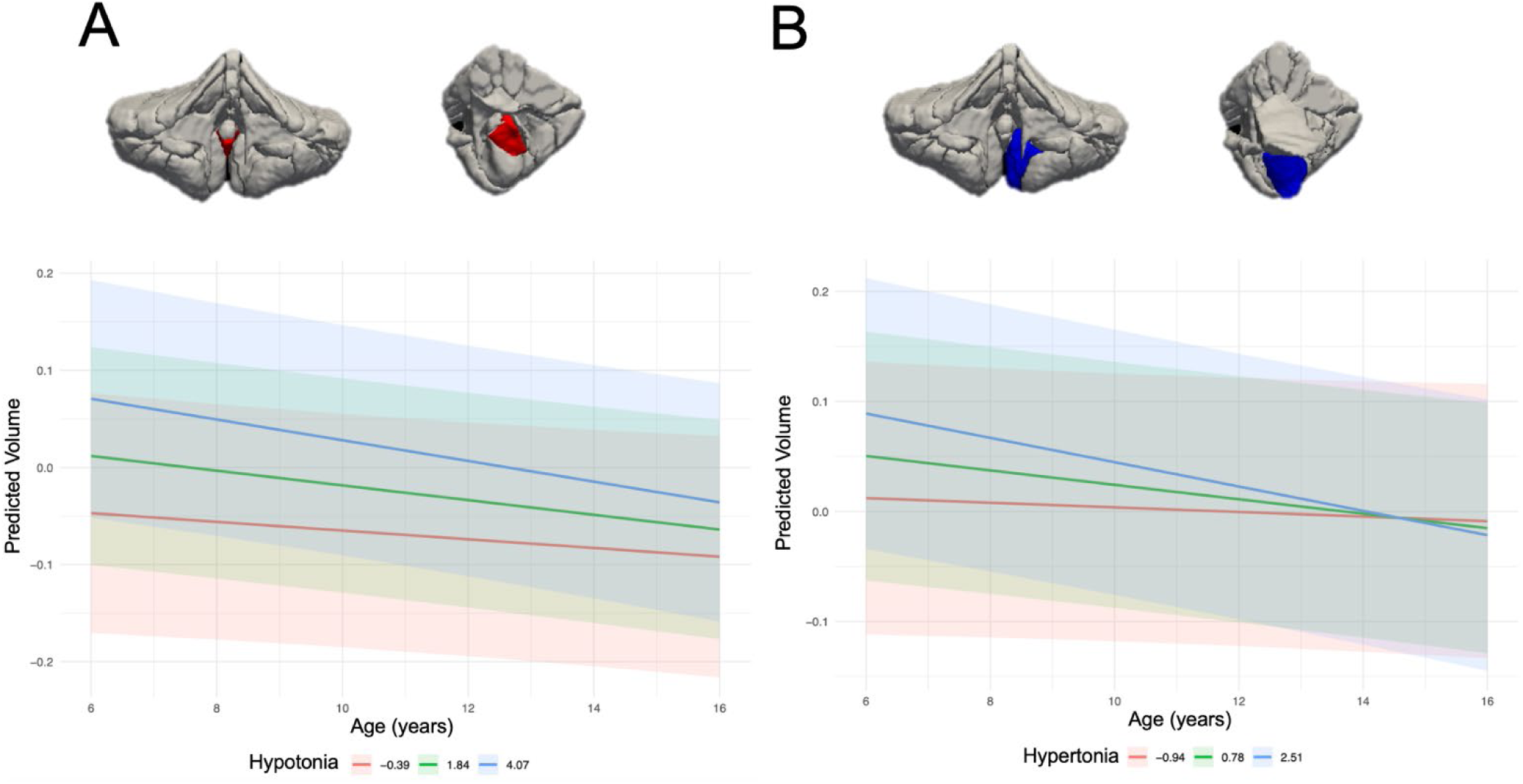
Anatomical Segmentations and Longitudinal Cerebellar Volume Trajectories by Infant Muscle Tone Severity. (**A**) Main effect of infant hypotonia on vermis IX volume (highlighted in red). (**B**) Interaction effect of infant hypertonia by age on left lobule IX volume (highlighted in blue). For each panel, top pictures show coronal and sagittal cerebellar views, while bottom plots show predicted volume trajectories across low, mean, and high severity scores of infant muscle tone. Lines represent mean trajectories, and shaded bands denote 95% confidence intervals.

**Figure 5.**
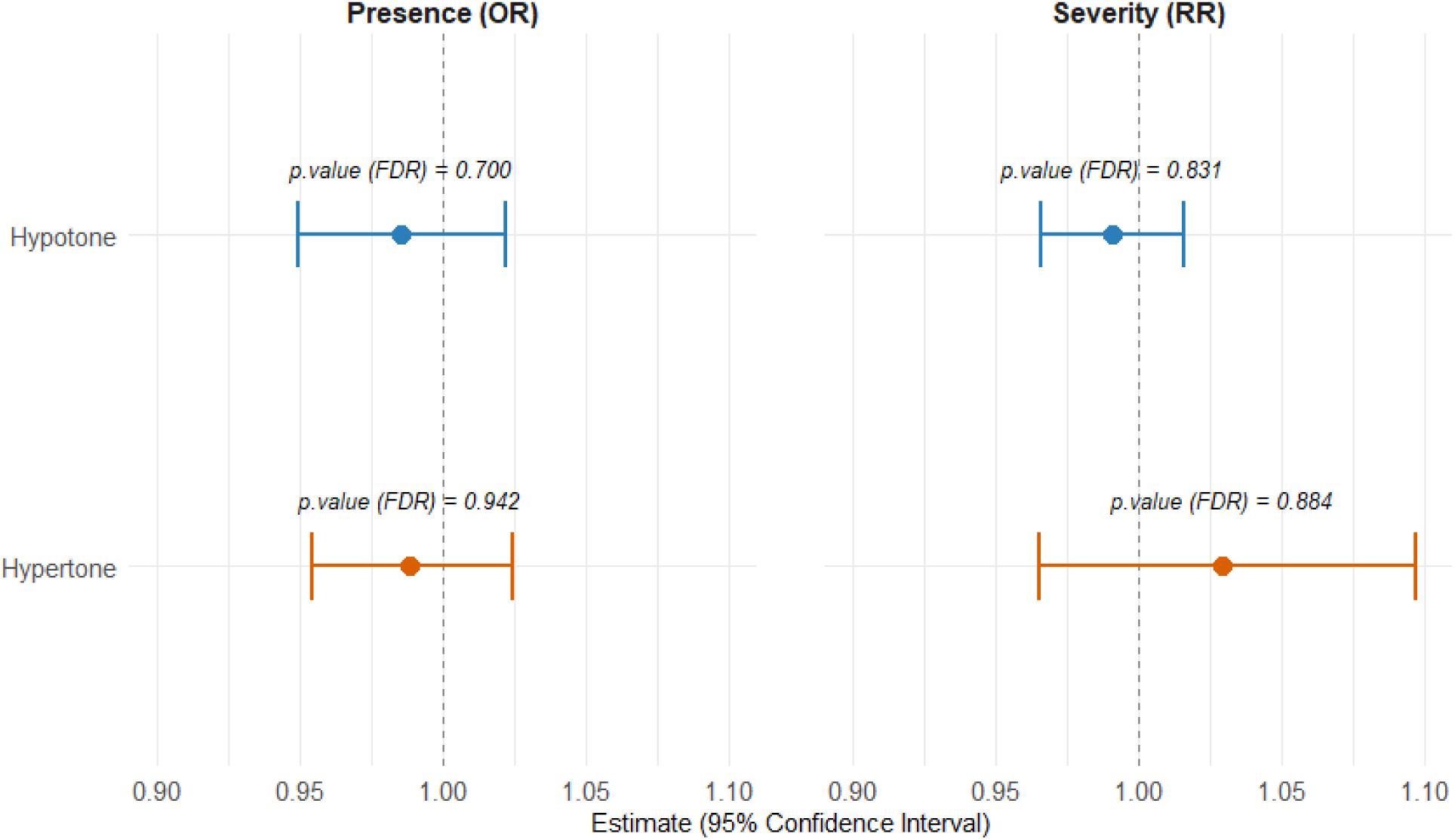
Associations Between Third-Trimester Fetal Transcerebellar Diameter and Infant Muscle Tone. *Note.* Forest plot of Hurdle regression estimates for third-trimester TCD predicting infant hypotonia (blue) and hypertonia (orange). Displayed are Odds Ratios (presence) and Relative Risks (severity) with 95% CIs. Dashed line indicates null effect (1.00). Models are adjusted for full prenatal covariate set; p-values are FDR-corrected.

**Table 3.** Summary of Longitudinal Main and Interaction Effects for Cerebellar Regions.

| *Hypotonia* |  |  |  |  |  |  |
| --- | --- | --- | --- | --- | --- | --- |
| ROI | Term | Estimate ($\beta$) | SE | 95% CI | *p* | *pFDR* |
| **Vermis IX** | **Main effect** | **0.037** | **0.015** | **[0.008, 0.066]** | **0.013** | **0.043** |
| Left Lobule VIIIB | Main effect | 0.040 | 0.019 | [0.001, 0.088] | 0.044 | 0.129 |
| Vermis VI | Main effect | 0.037 | 0.017 | [0.004, 0.069] | 0.027 | 0.084 |
| Vermis X | Main effect | 0.044 | 0.019 | [0.007, 0.082] | 0.021 | 0.079 |
| Vermis IX | Interaction effect | 0.002 | 0.001 | [0.004, 0.001] | 0.066 | 0.182 |
| Vermis X | Interaction effect | 0.003 | 0.002 | [0.006, 0.001] | 0.064 | 0.178 |
| Hand movement (left) | Interaction effect | 0.002 | 0.001 | [0.001, 0.005] | 0.081 | 0.213 |
| *Hypertonia* |  |  |  |  |  |  |
| Narrative | Main effect | 0.003 | 0.002 | [-0.007, 0.001] | 0.079 | 0.251 |
| Left Lobule IX | Main effect | 0.041 | 0.018 | [0.006, 0.076] | 0.021 | 0.652 |
| **Left Lobule IX** | **Interaction effect** | **0.003** | **0.001** | **[-0.005, -0.001]** | **0.012** | **0.041** |

*Note.* Bold values indicate significance at $pFDR < 0.05$ . Models are adjusted for sex, gestational age, ICV, age at MRI assessment, parental national origin, household income, maternal education, third-trimester maternal blood pressure, and prenatal substance use.

#### Infant Hypertonia

No structural or functional cerebellar ROIs demonstrated statistically significant main effects of hypertonia after FDR correction (Table 3). However, a statistically significant hypertonia-by-age interaction in left lobule IX was observed (*β*=−0.0027, *SE*=0.0011, *p*=0.012, *pFDR*=0.041, *95% CI* [−0.0048, −0.0006]; Table 3, Figure 4B). Individuals with typical muscle tone showed excepted flattened trajectory, whereas participants with elevated hypertonia scores demonstrated a declining trajectory across childhood and adolescence (Figure 4B). Additionally, no significant associations were observed across the functionally defined ROI’s after FDR correction (*pFDR*>0.05; supplement figure 3 and 4)

### Prenatal Transcerebellar Diameter and Infant Muscle Tone

Third-trimester fetal TCD was not significantly associated with subsequent infant muscle tone alterations after FDR adjustment (Figure 6). In hurdle regression models for hypotonia, TCD predicted neither symptom presence (*OR*=0.99, *95% CI* [0.95,1.02], *pFDR* =0.700) nor symptom severity among affected infants (*RR*=0.99, *95% CI* [0.97,1.02], *pFDR* =0.831). Similarly, no significant fetal associations were observed for hypertonia regarding either symptom presence (*OR*=0.99, *95% CI* [0.95,1.02], *pFDR* =0.942) or severity (*RR*=1.03, 95% *CI* [0.97,1.10], *pFDR* =0.884). Fetal cerebellar size during the third trimester does not appear to predict postnatally observed muscle tone variations.

### Cerebellar Volume as a Potential Mediator of Autistic Traits

To test whether structural variation in vermis IX mediates the link between infant hypotonia and adolescent autistic traits at age 14, causal mediation analyses were performed. In the total sample, vermis IX-volume did not mediate this association (ACME: *β*=−0.0004, *95% CI* [−0.0014,0.0006], *p*=.460; Supplemental Table 2). Neither the Average Direct Effect (ADE: *β*=0.0111, *95% CI* [−0.0060,0.0280], *p*=.203) nor the Total Effect (*β*=0.0107, *95% CI* [−0.0064,0.0277], *p*=.219) reached statistical significance in the pooled sample.

However, sex-stratified models revealed distinct patterns across groups. In boys, all pathways were non-significant, showing no indirect effect (*β*=−0.0006, *95% CI* [−0.0027,0.0015], *p*=.570) or direct effect (*β*=−0.0111, *95% CI* [−0.0384,0.0161], *p*=.422; Supplemental Table 3). In girls, while indirect mediation through vermis IX-volume was similarly absent (*β*=−0.0001, *95% CI* [−0.0013,0.0011], *p*=.853), a statistically significant direct effect was found (*β*=0.0256, *95% CI* [0.0041,0.0471], *p*=.020). This drove a significant total effect of infant hypotonia on adolescent SRS-scores among girls (*β*=0.0255, *95% CI* [0.0040,0.0470], *p*=.020; Supplemental Table 4). Thus, while vermis IX-volume does not function as a structural mediator, infant hypotonia demonstrates a direct, sex-specific association with adolescent autistic traits in females.

## Discussion

This study identifies three main insights into early cerebellar development, infant neuromotor signs, and adolescent autistic traits using a large, prospective population-based cohort. First, infant muscle tone alterations were associated with volume differences in cerebellar lobule IX. Infant hypotonia predicted volumetric differences in vermal lobule IX, whereas hypertonia showed age-dependent association with the left hemispheric lobule IX. Second, third-trimester fetal TCD did not predict infant muscle tone alterations in our study. Last, while vermis IX volume does not mediate the effect of infant muscle tone on adolescent autistic traits, infant hypotonia showed a small, direct association with higher autistic traits specifically in females. By utilizing a large, prospective birth cohort, we aimed to understand the cerebellum’s diverse contributions to early neurodevelopment and evaluate whether infant muscle tone may serve as an accessible early clinical marker of cerebellar differences and long-term neurodevelopmental risk.

Assessing infant muscle tone at 2–5 months of age captures a critical neurodevelopmental window which is also characterized by rapid postnatal expansion of cerebellar circuitry (Choe et al., 2013). Rather than evaluating isolated motor skills, muscle tone at this early age serves as a non-invasive clinical marker for the functional state and developmental pace of underlying neural circuits. In the present study, hypotonia and hypertonia were associated with vermal and hemispheric regions of lobule IX. Developmental functional mapping in infants implicated vermal and hemispheric parts of lobule IX to the cerebellum’s primary sensorimotor network (Lyu et al., 2024). Although both regions therefore share a broad sensorimotor classification, the cerebellar vermis and hemispheres functional and structural organization differs. The vermis, also called spinocerebellum, controls posture, core body muscles and balance, whereas cerebellar hemispheres are more involved in the coordination of fine distal movements and, in the case of posterior regions like lobule IX, are also connected to higher-order association networks including the default mode and frontoparietal networks in later development. (Kandel et al., 2021; Coffman et al., 2011; Amore et al., 2021). The localization to vermal lobule IX specifically, and to the left hemispheric lobule IX rather than other hemispheric sensorimotor regions such as lobules IV–VI and VIII, may reflect the specific developmental stage and clinical phenotype examined. Whereas lobules IV–VI and VIII are commonly associated with overt, goal-directed movement, infant muscle tone may more directly reflect tonic postural regulation, the ongoing background adjustments that maintain posture rather than discrete motor acts (Stoodley & Schmahmann, 2009; Guell et al., 2018). However, given our investigation in a large population sample, these regional findings should be interpreted with caution since effect sizes are small.

To test the directionality of our associations between infant muscle tone and later cerebellar volumes in lobule IX, we used third-trimester fetal TCD available in our population cohort. Fetal TCD was not associated with infant muscle tone which may reflect a true absence of association but likely reflects the limitations of transcerebellar diameter as a proxy for the localized subregional development relevant to muscle tone as well. A single one-dimensional ultrasound measurement lacks the spatial sensitivity to capture localized, subregional, or asymmetric tissue expansion (Scott et al., 2012). TCD measurements are also sensitive to fetal and maternal positioning during the acquisition of the ultrasounds, variability in acquisition protocol and operator technique across clinical sites, variability between operators and differences in ultrasound equipment between machines, all of which introduce measurement error that further limits its precision as a proxy for later, spatially specific cerebellar development (Volpe, 2009). These limitations suggest that third-trimester TCD is too crude a measure to reliably capture the localized cerebellar development potentially relevant to infant muscle tone.

We further examined whether vermal IX volume mediated the association between infant hypotonia and later autistic traits. Infant hypotonia showed a small direct association with adolescent autistic traits in females (*β*=0.0255), but not in males. Given the very small effect size, this finding should be interpreted cautiously and is more consistent with hypotonia serving as a subtle early vulnerability marker than as a major determinant of later autistic traits in our population sample. Sex-related differences in cerebellar maturation, motor development, and the expression or measurement of autistic traits could potentially contribute to this pattern (Lai et al., 2015; Werling & Geschwind, 2013). Replication in independent prospective cohorts, as well as in clinical samples is needed to determine whether this female-specific association reflects a robust association.

Strengths of this study include its prospective population-based design spanning 14 years, the use of detailed functional and anatomical cerebellar segmentations, normative modeling taking into account scanner differences, and Inverse Probability Weighting to handle attrition. Despite these methodological strengths, several limitations should be considered when interpreting our results. First, our neuroimaging analyses focused exclusively on the cerebellum, neglecting other subcortical and neocortical structures. This was a deliberate, hypothesis-driven choice, given that the cerebellum is consistently implicated in both motor tone regulation and autism across the structural neuroimaging literature (Stoodley & Schmahmann, 2009; Fatemi et al., 2012; Stoodley et al., 2017). As a result, we cannot draw conclusions about whole-brain structural associations, though restricting our analyses to the cerebellum also helped limit the multiple comparison burden given the large number of subregions already tested. Second, infant muscle tone was assessed at a single, potentially state-dependent time point, and hypotonia and hypertonia are clinically heterogeneous phenotypes that can arise from diverse underlying mechanisms and vary in severity. This heterogeneity introduces non-differential misclassification and measurement noise, which creates attenuation bias toward the null. Consequently, our population-based models may understate the true strength or anatomical specificity of these associations, or mask effects that are prominent only within specific clinical sub-phenotypes. Third, the available MRI resolution (1mm isotropic) limited the precise delineation of narrow vermal structures, which is particularly relevant for our vermal lobule IX findings given that the vermis is a small, midline structure where partial volume effects and segmentation errors are most likely to affect volume estimates. Finally, the study used continuous SRS traits rather than diagnostic outcomes.

Consequently, these findings speak to population-level variation in autistic traits and may not directly generalize to individuals with a clinical ASD diagnosis, where symptom severity and co-occurring conditions could alter these associations. Similarly, the sex-specific mediation findings may be susceptible to limited statistical power, as stratifying by sex substantially reduced the sample size within each group, and mediation models require larger samples than simple bivariate associations to reliably detect indirect effects.

Future research should adopt a whole-brain, multimodal approach potentially combining cerebellar and cerebral morphometry with diffusion MRI and resting-state functional connectivity. Such studies could clarify whether early differences in vermal IX and the left hemispheric lobule IX reflect isolated cerebellar variation or atypical development of broader cerebro-cerebellar networks. Repeated assessments of infant muscle tone and motor behavior are also needed to distinguish persistent neuromotor characteristics from state-dependent findings. Larger cohorts with repeated fetal, infant, and childhood imaging, formal tests of sex moderation, and longitudinal assessments of autistic traits will be important for determining the developmental and clinical significance of the observed associations. Finally, replicating these associations in clinical samples with neuromotor or neurodevelopmental difficulties would help establish whether infant muscle tone can serve as a marker in populations where early identification is most clinically relevant.

In conclusion, early infant muscle tone deviations were associated with localized differences in cerebellar lobule IX, with hypotonia and hypertonia mapping onto functionally distinct vermal and hemispheric subregions respectively. Global prenatal measurements such as fetal transcerebellar diameter did not reflect these localized postnatal alterations, possibly because available ultrasound measurements were not sensitive enough to detect associations with localized postnatal cerebellar development. Furthermore, while infant hypotonia displayed a small, direct longitudinal association with adolescent autistic traits in females, vermis IX volume did not mediate this pathway. Future research should examine what other mechanisms may explain this association. Together, these findings implicate early muscle tone evaluations as a meaningful marker of subregional cerebellar organization, while underscoring that it captures only part of the neurodevelopmental pathway to autistic traits.

