## Supplementary Materials for "Neonatal Muscle Tone Predicts Cerebellar Morphology Later in Development Without Mediating Autistic Traits"

**Supplementary Table 1**
*Items on Neurological Assessment of Infant Muscle Tone.*

| Subscale | Item Description | Optimal | Non-optimal (Hypotonia) | Non-optimal (Hypertonia) |
| --- | --- | --- | --- | --- |
| Supine | Resting posture | Semi-flexed legs; slight abduction at the hips | Legs flat on the surface | Legs stretched |
|  | Adductor angle | >80∘ to <140∘ | >140∘ | <80∘ |
|  | Popliteal angle | 90∘ to 130∘ | 130∘ to 180∘ | <90∘ |
|  | Ankle angle | >20∘ to <90∘ | <20∘ | >90∘ |
|  | Head preference | No | Yes | — |
|  | Opening & closing hands | Yes | Sometimes closed | Always closed |
|  | Alternating leg movements | Yes | Decreased | Absent |
|  | Grasps with one hand | Yes | Decreased | Absent |
|  | Hyperextension | No | Sometimes | Yes |
|  | Dyskinesia | No | Sometimes | Yes |
| Supine-to-sit | Traction response | Arms moderately flexed | Arms fully extended, no resistance | Strong resistance, flexion elbows, legs extended |
|  | Traction response - head control | Active lift of head | Head lag | Exaggerated |
| Horizontal | Ventral Tone | Normal tone | Low tone | Back and limbs stretched |
| Vertical | Head | Normal tone | Low tone | High tone |
|  | Shoulders | Normal tone | Low tone | High tone |
|  | Trunk | Normal tone | Low tone | High tone |
|  | Legs | Normal tone | Low tone | High tone |
| Prone | Pulls arms up | Yes | No |  |
|  | Turns head | Yes | No |  |
|  | Lifts head | Yes | No | Overstretched |
| Sitting | Needs support | Yes | No |  |
|  | Head control | Yes | No |  |
|  | Shoulder retraction | No | Yes |  |
|  | Shape of the back | Round | Straight | Scoliosis |

| Effect | *Estimate* | *SE* | 95% CI | *p* |
| --- | --- | --- | --- | --- |
| ACME (Indirect Effect) | -0.0004 | 0.0005 | [-0.0014, 0.0006] | .459 |
| ADE (Direct Effect) | 0.0111 | 0.0087 | [-0.0060, 0.0280] | .203 |
| Total Effect | 0.0106 | 0.0087 | [-0.0064, 0.0277] | .219 |
| Proportion Mediated | -0.0200 | 0.2094 | [-0.4304, 0.3903] | .923 |
| *Note. ACME =* Average Causal Mediation Effect*; ADE =* Average Direct Effect*; CI =* confidence interval*.* | | | | |

**Supplementary Table 2**
*Final Pooled Mediation Analysis Results: Hypotonia and Cerebellar Middle Vermis Lobule IX.*

**Supplementary Table 3**
*Final Pooled Mediation Analysis Results: Hypotonia and Cerebellar Middle Vermis Lobule IX for Boys.*

| Effect | *Estimate* | *SE* | 95% CI | *p* |
| --- | --- | --- | --- | --- |
| ACME (Indirect Effect) | -0.0006 | 0.0011 | [-0.0027, 0.0015] | .570 |
| ADE (Direct Effect) | -0.0111 | 0.0139 | [-0.0384, 0.0161] | .422 |
| Total Effect | 0.0118 | 0.0139 | [-0.0391, 0.0156] | .399 |
| Proportion Mediated | 0.0171 | 0.3353 | [-0.6401, 0.6742] | .959 |
| *Note. ACME =* Average Causal Mediation Effect*; ADE =* Average Direct Effect*; CI =* confidence interval*.* | | | | |

| Effect | *Estimate* | *SE* | 95% CI | *p* |
| --- | --- | --- | --- | --- |
| ACME (Indirect Effect) | -0.0001 | 0.0006 | [-0.0013, 0.0011] | .853 |
| ADE (Direct Effect) | 0.0256 | 0.0110 | [0.0041, 0.0471] | .020 |
| Total Effect | 0.0255 | 0.0110 | [0.0040, 0.0470] | .020 |
| Proportion Mediated | -0.0015 | 0.0383 | [-0.0745, 0.0761] | .968 |
| *Note. ACME =* Average Causal Mediation Effect*; ADE =* Average Direct Effect*; CI =* confidence interval*.* | | | | |

**Supplementary Table 4**
*Final Pooled Mediation Analysis Results: Hypotonia and Cerebellar Middle Vermis Lobule IX for Girls.*

**Supplementary Figure 1**

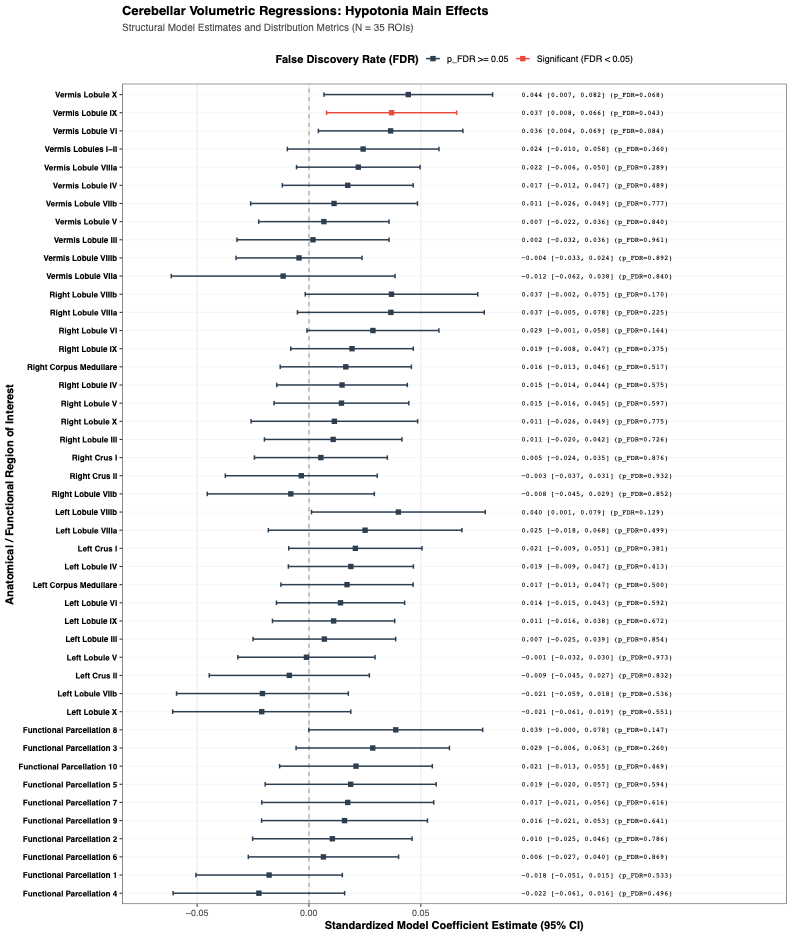

**Supplementary Figure 2**

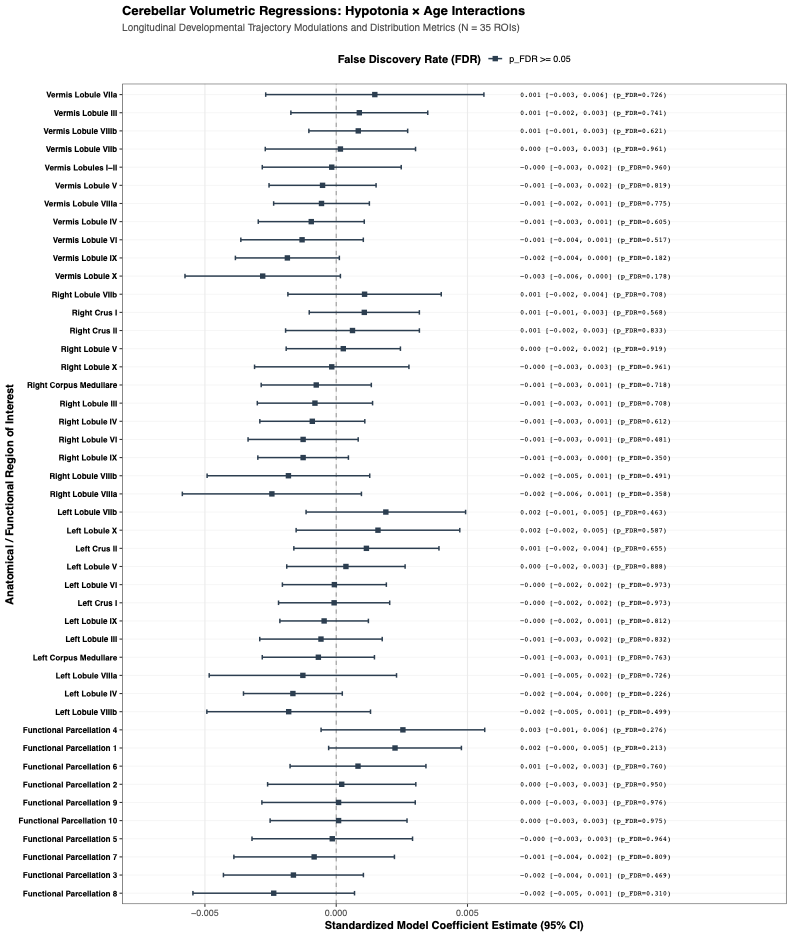

**Supplementary Figure 3**
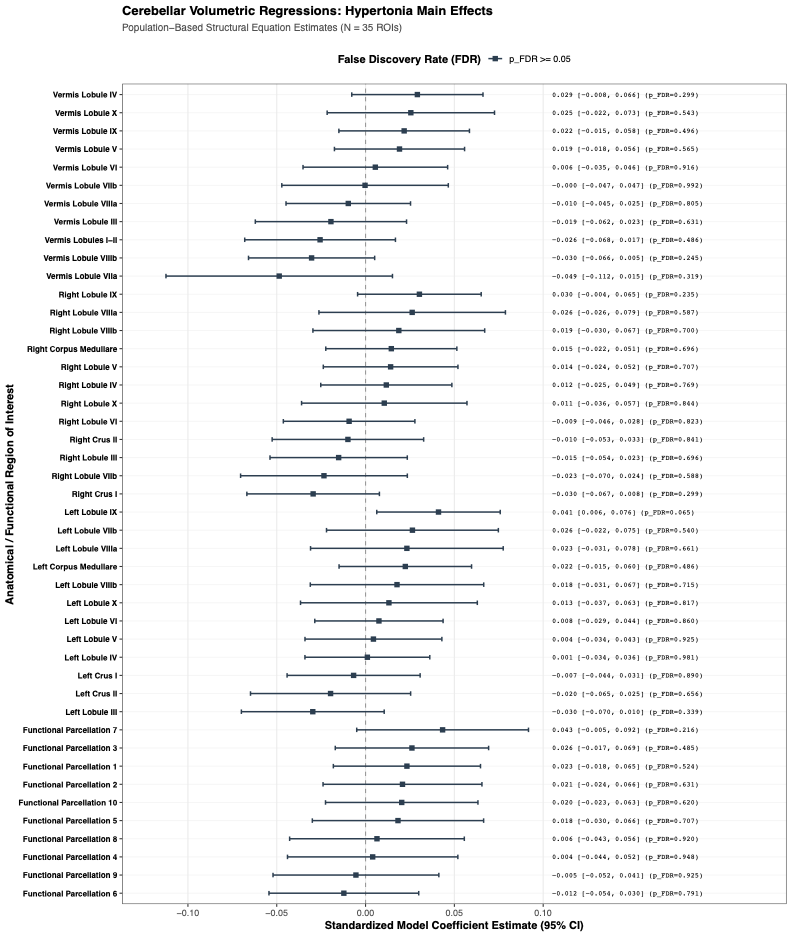

**Supplementary Figure 4**

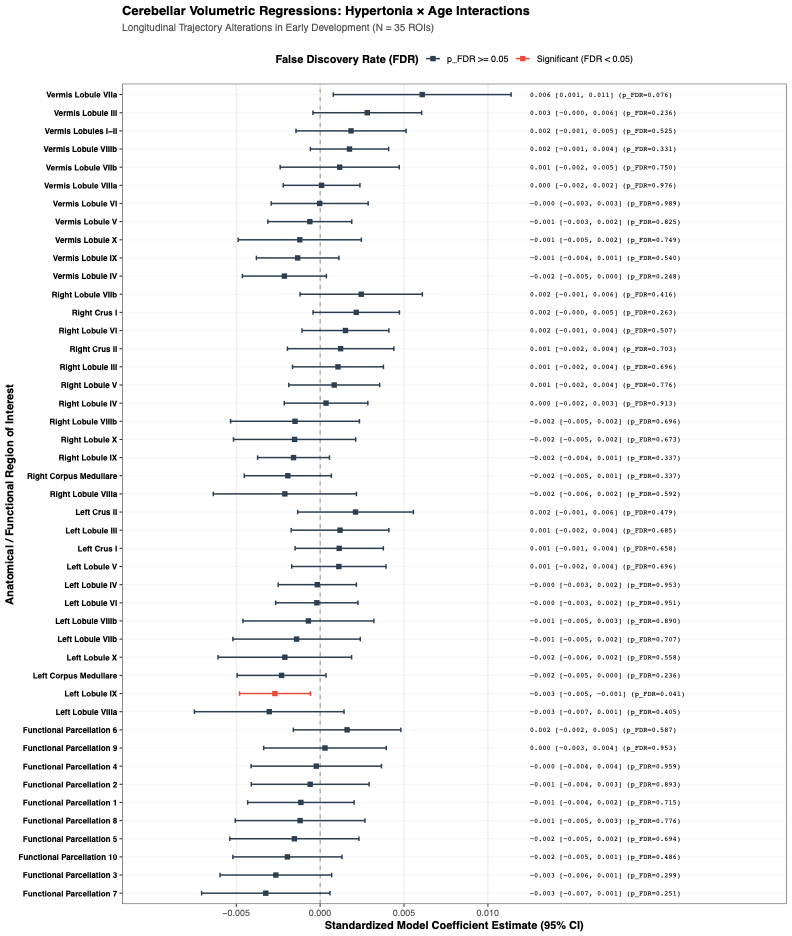
